# Internally-consistent and fully-unbiased multimodal MRI brain template construction from UK Biobank: Oxford-MM

**DOI:** 10.1101/2023.11.30.569378

**Authors:** Christoph Arthofer, Stephen M. Smith, Gwenaëlle Douaud, Andreas Bartsch, Fidel Alfaro-Almagro, Jesper Andersson, Frederik J. Lange

## Abstract

Anatomical MRI templates of the brain are essential to group-level analyses and image processing pipelines, as they provide a reference space for spatial normalisation. While it has become common for studies to acquire multimodal MRI data, many templates are still limited to one type of modality, usually either scalar or tensor-based. Aligning each modality in isolation does not take full advantage of the available complementary information, such as strong contrast between tissue types in structural images, or axonal organisation in the white matter in diffusion tensor images. Most existing strategies for multimodal template construction either do not use all modalities of interest to inform the template construction process, or do not use them in a unified framework.

Here, we present multimodal, cross-sectional templates constructed from UK Biobank data: the OMM-1 template, and age-dependent templates for each year of life between 45 to 81. All templates are fully unbiased to represent the average shape of the populations they were constructed from, and internally consistent through jointly informing the template construction process with T1, T2-FLAIR and DTI data. The OMM-1 template was constructed with a multi-resolution, iterative approach using 240 individuals in the 50-55 year age range. The age-dependent templates were estimated using a Gaussian Process, which describes the change in average brain shape with age in 37,330 individuals.

All templates show excellent contrast and alignment within and between modalities. The global brain shape and size is not preconditioned on existing templates, although maximal possible compatibility with MNI-152 space was maintained through rigid alignment. We showed benefits in registration accuracy across two datasets (UK Biobank and HCP), when using the OMM-1 as the template compared with FSL’s MNI-152 template, and found that the use of age-dependent templates further improved accuracy to a small but detectable extent. All templates are publicly available and can be used as a new reference space for uni- or multimodal spatial alignment.

## 1. Introduction

Anatomical magnetic resonance imaging (MRI) templates of the brain aim to provide representative models of average shape and voxel signal intensity of the populations from which they were constructed. They are essential for many different kind of neuroimaging analyses as they provide a common reference space for the spatial normalization of individual subjects using image registration methods. The resulting transformations and derived measures, such as Jacobian determinant maps, between each individual and a template, and the transformed images in template space, enable the study of intra- and inter-group variability or agreement, unbiased group comparisons of within-subject longitudinal changes and atlas-based segmentation of regions of interest (ROIs) at subject level.

Template construction methods aim to find an average intensity and average shape template, i.e., the shape and intensity of the template should, on average, not be more like any one individual than any other (see section 2.3.2 for mathematical description). This is typically achieved through a series of steps to avoid bias in appearance or shape towards any single individual. The most commonly used method is based on an iterative framework (Guimond et al., 1998, 2000), which was later extended into a multi-resolution approach with a hierarchical processing structure (Grabner et al., 2006; Fonov et al., 2011). First, individual images are corrected for global (affine) misalignment using translation, rotation, scale and shear, which allows for the construction of an initial average affine template. Each individual is then iteratively non-linearly registered to the current template (starting with the affine template in the first iteration), followed by spatial unbiasing of the warps, and resampling of the subject images. Finally, the average across the resampled images becomes the new template and serves as the reference space for the next iteration. These steps are repeated until convergence, while warp resolution and image blurring are adjusted from coarse to fine.

Existing templates are often described as uni- or multimodal based on the number of modalities they comprise. An overview of some of the most commonly used and some more recent templates can be found in Table 1. In contrast to one modality in unimodal templates, multimodal templates aim to provide volumes of different, but anatomically-corresponding, modalities. This notion of multimodality in most existing templates stems from the post hoc availability of multiple modalities in template space, but generally does not refer to the modalities used during the template construction process. Driving this process with complementary information from different modalities of interest is highly desirable since it can improve registration quality. For example, the axonal organisation derived from diffusion imaging data can add valuable information about the white matter, which would not be available from T1-weighted (T1) images only. Some existing templating methods use one modality to drive the construction, e.g., T1, and then apply the same deformation fields to all modalities of interest, e.g., T2-weighted (T2) or diffusion tensor images (DTI) (Rohlfing et al., 2010; Fonov et al., 2011; Gupta et al., 2016). Others use modalities *derived from* the modality of interest, e.g., fractional anisotropy (FA) maps from DTI, to drive the construction and then transform the modality of interest (DTI) with the same transformations (Zhang et al., 2011; Lv et al., 2022). Estimating deformation fields based on a subset of modalities or surrogates, and applying the same deformation fields to all other modalities is not optimal. This strategy can lead to unwanted biasing effects in the template, since not all modalities contribute to the estimation of the deformation fields that are used for resampling and spatial unbiasing. This might not have a large impact when using modalities with similar information content, for example, when estimating a warp based on T1 images and applying the same warp to T1 and T2 images. However, for modalities with different information content it could introduce a spatial bias. For example, estimating deformation fields based on structural or diffusion-derived scalar modalities, and applying them to diffusion tensors could lead to a bias in the location or orientation of the diffusion data.

**Table 1:**
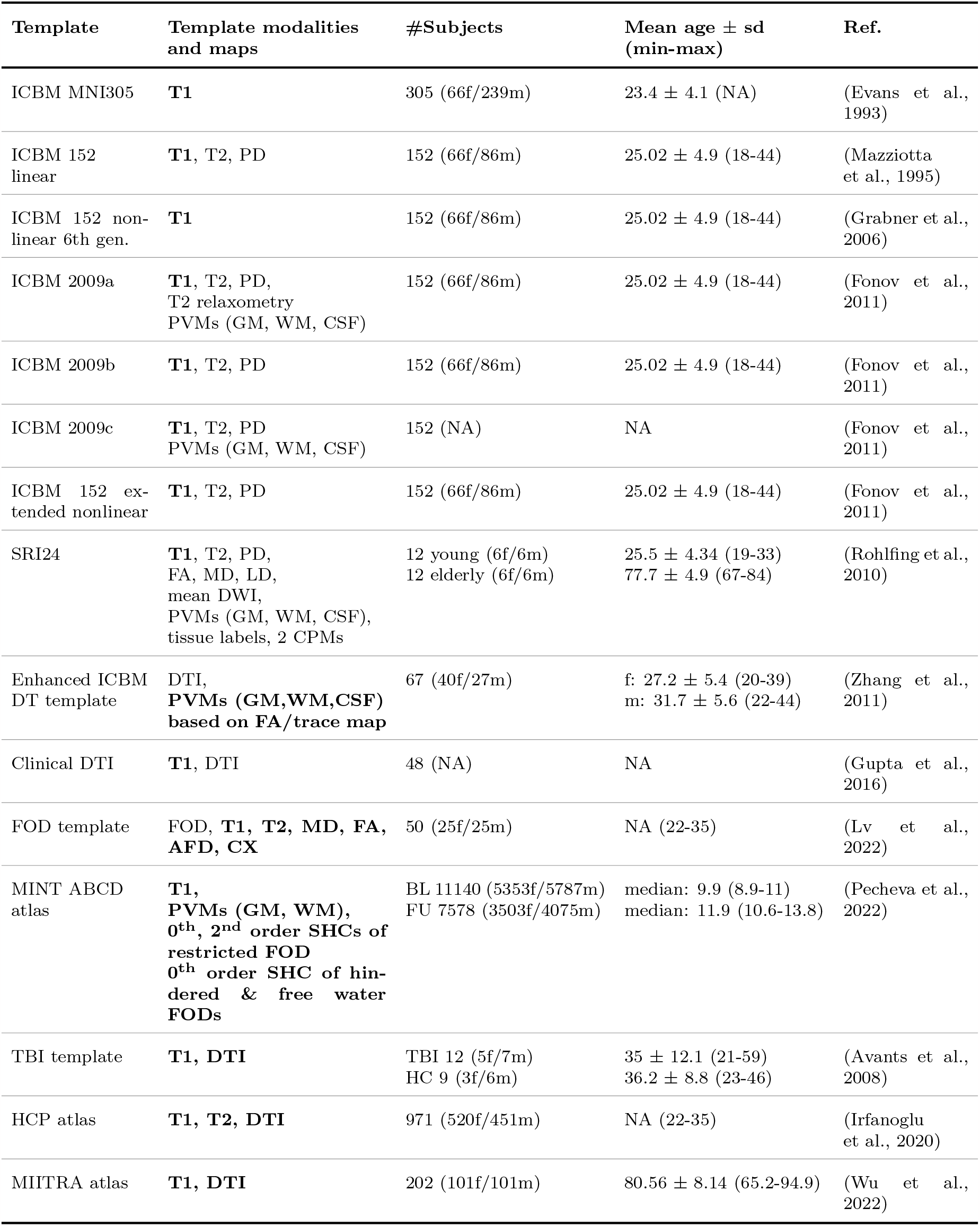
Overview of existing unimodal and multimodal templates. **Modalities in bold** are used in the construction. PD…proton-density weighted, LD…longitudinal diffusivity, DWI…diffusion-weighted imaging, PVMs…partial volume maps, CPMs…cortical parcellation maps, SHC…spherical harmonics coefficient, AFD…apparent fibre density, CX…fibre complexity, FOD…fibre orientation distribution, BL…baseline, FU…follow up

One fully-unbiased multimodal (FUMM) template was constructed from individuals in the adolescent brain and cognitive development (ABCD) study. For this template, eleven scalar modalities, including three structural modalities and eight dMRI-derived modalities but no DTI data were used as in-put to the Multimodal Image Normalisation Tool (MINT) (Pecheva et al., 2022). Another FUMM templating strategy for scalar and tensor modalities was applied in the construction of the MIITRA template (Wu et al., 2022). The method alternates between registrations within each of the T1 and DTI modalities. In each iteration, deformation fields are estimated within one of the two modalities with a modality-specific registration method. The same transformations are applied to data from both modalities in all iterations except the last, where the DTI data undergoes one more transformation that is not applied to the T1 images. A similar iterative approach, involving multiple repeated registrations with the two methods, is required when spatially normalising individuals to the MIITRA template. Since both modalities drive the template construction the resulting templates are fully unbiased. However, the use of two methods does not provide a unified and internally-consistent framework. To the best of our knowledge, the only two methods that can accommodate both scalar and tensor modalities, and, consequently, allow fully-unbiased and internally-consistent template construction, are Symmetric Normalization for Multivariate Neuroanatomy (SyNMN) (Avants et al., 2008) and DR-TAMAS (Irfanoglu et al., 2016). SyNMN was applied in the construction of a combined T1 and DTI template to investigate traumatic brain injury (TBI) and later in the construction of a template from arterial spin labelling, T2-weighted-Fluid-attenuated inversion recovery (T2-FLAIR), DTI, functional MRI (fMRI), T1 and T2 data (Tustison et al., 2015). The SyNMN tool and templates are not publicly available at the time of writing. DR-TAMAS has been used for the construction of a DTI atlas (Irfanoglu et al., 2020) from the Human Connectome Project Young Adult (Van Essen et al., 2012) dataset (22 - 35 year age range). This DTI template also comprises T1 and T2 volumes, and all modalities were used to drive the registrations during the template construction process. The atlas was constructed from 971 individuals and has good levels of detail and contrast (although not quite as good as might be hoped for, given the quality of the data and the number of subjects).

Most existing multimodal templates provide a single, cross-sectional average of brain shape and intensity from the subjects in a cohort. However, arguably, a template should also be similar to a given population under investigation to reduce the amount of deformation required when aligning individuals to it. The main factor contributing to morphological variability in large datasets is the subjects’ age range. As datasets become larger in size and the subjects’ age range within datasets increases, it becomes more difficult to capture the age-related increase in brain shape variability in a single template. Spatiotemporal, or age-dependent templates (ADTs), for sub-populations with smaller age ranges can provide more similar reference spaces. Several construction methods based on discrete bins (Fillmore et al., 2015), kernel regression (Davis et al., 2007; Serag et al., 2012) and neural network-based architectures (Dalca et al., 2019; Xia et al., 2019; Zhao et al., 2019; Wilms et al., 2020) have been described in the literature. However, these have been mainly used for unimodal ADT construction and, to the best of our knowledge, do not publicly provide multimodal templates with scalar and tensor modalities for general use.

### 1.1. Summary of our work

The main contributions of our work include the construction of a cross-sectional, internally consistent and fully-unbiased multimodal, whole-head template, the Oxford-MultiModal-1 (OMM-1), and the development of a modelling and prediction-based approach, which was applied in the construction of multimodal, average-shape ADTs.

The former was obtained from 240 UK Biobank (UKB) individuals (50-55 years, 50% females) with the iterative approach described in (Fonov et al., 2011). First, we constructed an unbiased affine template, which was refined from coarse to fine by iterating through nonlinear registrations, and unbiasing, warping and averaging steps. The ADTs were obtained by nonlinearly registering 37,330 UKB individuals (45-82 years) to OMM-1 and using the acquired deformation fields and corresponding individuals’ ages to model the change in average brain shape with age using a Gaussian Process (GP). Finally, the trained model allowed us to predict and apply a mean deformation field for each year of age to derive age-dependent templates from the initial 240 UKB individuals.

The OMM-1 and its associated ADTs provide anatomically-corresponding scalar (T1 and T2-FLAIR) and tensor (DTI) volumes. These same modalities were used to drive the construction process by simultaneously informing the nonlinear registrations. These registrations were performed with FSL’s MultiModal Registration Framework (MMORF) (Lange et al., 2020a,b), which estimates a single warp by optimizing over an arbitrary number of scalar and tensor input modalities. This ensures internal consistency by avoiding the need to use different registration methods for different modalities, and full-unbiasing of all volumes with respect to all modalities of interest. Our template construction pipeline provides a unified framework that can easily be extended or adjusted to other scalar or tensor modalities of interest. Since the OMM-1 is unbiased with respect to the UKB subjects from which it was created, its size and shape differs from the most commonly used template, the MNI 152 (in its various revisions), as MNI 152 templates are not very close to representing the size of the average adult brain. However, the OMM-1 was rigidly (6 degrees of freedom) aligned to MNI space, and transformations between the two templates are provided, to aid compatibility when switching between them. Finally, we have investigated the benefits of using our multimodal templating framework for spatial normalization in age-diverse populations of two datasets.

## Methods

### Data

In this work we used scalar- and tensor-valued, non-defaced brain MRI data from UKB (Miller et al., 2016), one of the largest prospective epidemiological studies to date, which aims to acquire multimodal MR imaging data from 100,000 participants.

Imaging data from three MRI modalities including T1, T2-FLAIR and DTI were used for template construction. T1 provides information about the basic anatomical structure of the brain and shows strong contrast between the main tissue classes (gray and white matter, and cerebrospinal fluid). It is acquired as part of most imaging studies and has become the core modality of choice for existing adult human templates. T2-FLAIR was included as a second structural modality due to its enhanced contrast of subcortical gray matter regions, such as the striatum, pallidum, substantia nigra, red nucleus and dentate nucleus, of the olfactory bulbs, but also between normal appearing white matter and white matter hyperintensities. Diffusion MRI provides information about the properties of the local tissue microstructure and white matter tract structure. It makes it possible to estimate a diffusion tensor for each voxel (Basser et al., 1994) that adds information about the axonal organisation and the preferred directions of diffusion. We decided to use non-defaced T1 and T2-FLAIR data to construct a template that is sharp and clear in both intra- and extracranial regions and, hence, may be useful for a variety of applications.

All imaging data were collected at one of three UKB sites using identical 3T Siemens Skyra scanners running VD13 and a standard Siemens 32-channel receive head coil. A brief overview of the parameters used to acquire T1, T2-FLAIR and dMRI can be found in Table 2. For a detailed description of the acquisition protocol in the UKB brain imaging study we refer the reader to (Miller et al., 2016).

**Table 2:** UK Biobank brain MRI acquisition parameters for T1, T2-FLAIR and dMRI from (Miller et al., 2016). R=in-plane acceleration factor, MB=multiband factor, PF=partial Fourier.

| Modality | Voxel size,<br>Matrix | Key parameters |
| --- | --- | --- |
| T1 | $1.0 \times 1.0 \times 1.0$ mm,<br>208 × 256 × 256 | 3D MPRAGE, sagittal, R=2, TI/TR=880/2000 ms |
| T2-FLAIR | $1.05 \times 1.0 \times 1.0$ mm,<br>192 × 256 × 256 | FLAIR, 3D SPACE, sagittal, R=2, PF 7/8, fat sat,<br>TI/TR=1800/5000 ms, elliptical |
| dMRI | $2.0 \times 2.0 \times 2.0$ mm,<br>104 × 104 × 72 | MB=3, R=1, TE/TR=92/3600 ms, PF 6/8, fat sat,<br>b=0 s/mm<sup>2</sup> (5× + 3× phase-encoding-reversed),<br>b=1000 s/mm<sup>2</sup> (50×)<br>b=2000 s/mm<sup>2</sup> (50×) |

The OMM-1 was constructed from 240 individuals uniformly and randomly sampled from the 50-55 year age range (40 individuals per year, 50% female). The size of the sample was informed by previous investigations on the Human Connectome Project dataset, where Yang et al. (Yang et al., 2020) have shown that sample sizes larger than 200 individuals are associated with only small changes to the final templates. We selected individuals from the younger end of the UKB age range that provided sufficient data for uniform sampling. This minimises the appearance of ageing-related features and, therefore, maximises the utility of the template when applied to studies involving younger subjects.

All UKB images went through the manual and automated quality control (QC) pipeline described in (Alfaro-Almagro et al., 2018). Although badly corrupted images are excluded by this pipeline, several additional criteria for subjects to be considered in our random sample were defined. These requirements included the availability of all three modalities, less than 0.5% of the total brain volume containing white matter hyperintensities, and small alignment discrepancies. Alignment discrepancy measurements had been calculated as the correlation ratio between registered within-subject modalities by the QC pipeline and are available as QC imaging-derived phenotypes (IDPs) for all three modalities. Extreme scores are potential indicators for poor alignment, or the presence of artefacts or outliers. In particular, we used IDPs that describe the discrepancies between an individual’s T1 structural image and the MNI 152 6th gen. (Grabner et al., 2006) after nonlinear alignment, and between the T2-FLAIR and the corresponding T1 image, and the dMRI and the corresponding T1 image after linear alignment. Thresholds of 0.5 for the T1 and T2-FLAIR and 0.6 for the dMRI discrepancies were applied, to allow for a large enough sample of subjects from the selected age groups.

For the construction of the age-dependent templates, images from 37,330 (age 45-82 years) individuals, which had T1, T2-FLAIR and dMRI data, were used. Given this large sample, the image quality at the individual level is expected to have less impact on the final average templates for age modelling compared to the smaller sample used for the OMM-1. Therefore, no further selection criteria were applied.

### 2.2. Data preprocessing

We used both minimally processed and preprocessed UKB imaging data. The T1, T2-FLAIR and dMRI volumes of the former are gradient-distortion corrected, and the T1 and T2-FLAIR volumes are not defaced, i.e., they include parts of the neck, nose and mouth. The latter had been preprocessed with the standard pipeline described in (Alfaro-Almagro et al., 2018), which, in addition to gradient-distortion correction, includes defacing, cropping, brain extraction through atlas-based mask propagation, and intensity inhomogeneity correction of T1 and T2-FLAIR images. Brain-extracted T2-FLAIR and dMRI images are rigidly co-registered to the corresponding individual’s T1 reference space using the B0s as the moving image and boundary-based registration (Greve and Fischl, 2009) as the cost function in FSL’s FLIRT (Jenkinson and Smith, 2001). dMRI data are corrected for susceptibility and eddy current distortion, as well as head motion, using FSL’s topup (Andersson et al., 2003) and eddy (Andersson and Sotiropoulos, 2016; Andersson et al., 2016) before fitting the diffusion tensors (Basser et al., 1994) on the b=1000 images (50 directions) with FSL’s DTIFIT^1^. This standard preprocessing pipeline was extended for the template construction pipeline. The binary brain masks in individual dMRI spaces were slightly reduced in size by smoothing with an un-normalised mean filter (3 x 3 x 3 kernel size, to create a smooth transition between brain and background), thresholding at 0.9, and eroding by one voxel. These masks were used to reduce the impact of noisy DTI voxels at the border of, and outside the brain during nonlinear registrations. Bias fields created with FSL’s FAST were transformed from each individual’s reference spaces to their non-defaced T1 and T2-FLAIR native spaces and used to correct for intensity inhomogeneity in the brain. High intensity values of the scalp in T1 images were smoothly-clamped with a custom function (see Appendix A) to avoid negative effects on the non-linear registrations during template construction. We did not perform any resampling with the transformations estimated between modalities to avoid the accumulation of interpolation errors.

In the rest of this manuscript, we will use the following notation: the set of *N* subjects, where each individual *n* has data from three modalities *M* is defined as

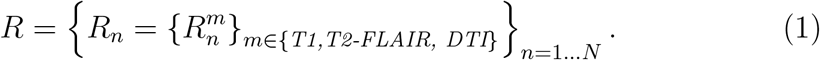

### 2.3. Template construction

Our multimodal template and age-dependent template construction pipeline consists of three main parts.

First, an unbiased affine template was constructed by correcting for global (affine) misalignment between individuals (Fig. 1A, Section 2.3.1), which was then rigidly aligned to MNI space (Grabner et al., 2006).

**Figure 1.**
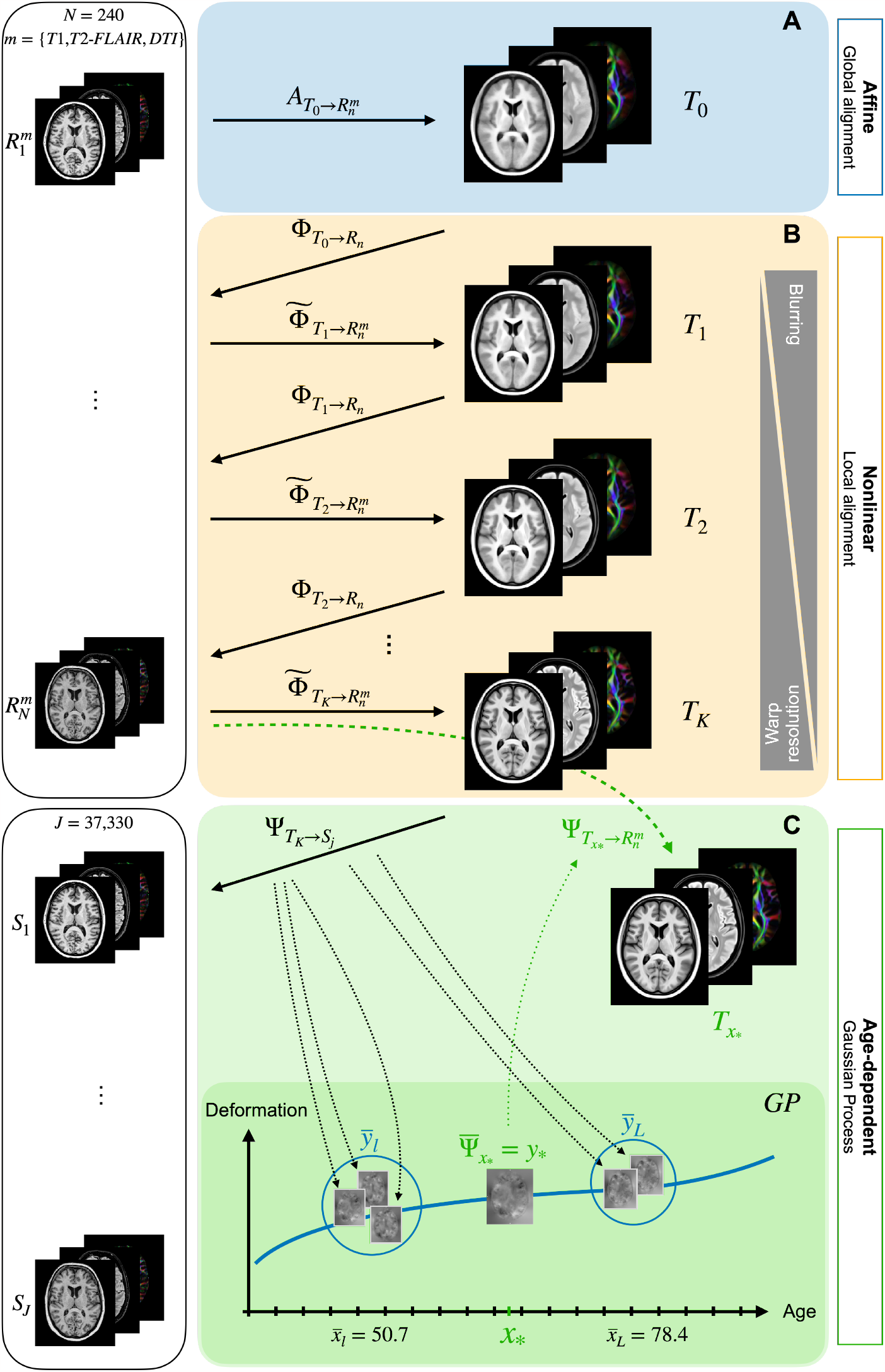
Scalar and tensor-based modalities from 240 UKB individuals were used to construct (A) the unbiased affine template by correcting for global misalignment and (B) the final nonlinear O MM-1 template by iterating through the hierarchical optimization approach. (C) Age-dependent templates were derived from the predictions of a Gaussian process model trained on ages, and warps to OMM-1 space from 37,330 UKB individuals.

Second, this affine template was used to initialise a nonlinear, hierarchical, multi-resolution templating approach (Fonov et al., 2011), which iterated through registration, unbiasing, transformation, and averaging steps (Fig. 1B, Section 2.3.2). The final nonlinear template, the OMM-1, represents the average shape and intensity of the 240 individuals on which it is based.

Third, the OMM-1 was used as a template to spatially normalise 37,330 individuals from the UKB imaging cohort, resulting in one deformation field for each subject (Fig. 1C, Section 2.3.3). A Gaussian process (GP) was used to model the morphological differences captured by these deformation fields as a function of age. After fitting the model, a mean deformation field was predicted for every year of age between 45-81, and used to generate the corresponding age-dependent template (ADT). In the following sections we will discuss each of these steps in detail.

#### 2.3.1. Affine template construction

An initial affine template was constructed from the preprocessed, brain-extracted *R*^*T1*^ images. One subject was randomly selected as a reference space and the remaining subjects were affinely registered to this reference with 12 degrees of freedom (DOF). To avoid the introduction of a bias towards the brain geometry of the reference individual, the transformation from each individual’s space to the mid-space of all subjects was calculated using FSL’s *midtrans* function. We performed preliminary tests with different subjects as an initial reference, and confirmed that we did not find any difference in the final results. Brain-extracted *R*^*T1*^ images were resampled into this unbiased space by applying the corresponding transformations to them. The first affine template was created by calculating voxel-wise the median over the resampled images, which provides a sharper group average compared with taking the mean at this early stage (Fig. 1A) and was found to improve registration performance in the subsequent iterations.

This initial template was rigidly (6 DOF) aligned to the space of the asymmetric version of the nonlinear 6th gen. ICBM 152 template (MNI 152) (Grabner et al., 2006) included in FSL, to maximise similarity between the spaces while avoiding shearing and scaling effects. The final set of linear transformations 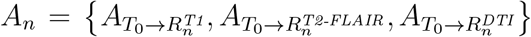for each individual were created by concatenating the corresponding rigid transformation from each modality’s native space to T1 reference space, the affine transformation from the T1 reference space to the unbiased template space, and the rigid transformation to the space of the new template *T*_0_. Non-defaced images were transformed from their native spaces to *T*_0_ by applying the corresponding concatenated transformations using spline interpolation. Additionally, T1 and DTI brain masks were resampled with the same transformations using trilinear interpolation.

The voxel-wise median of the resampled images for each of the modalities provided the final affine template with three volumes 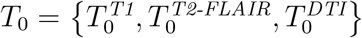.Similarly, mean T1 and DTI brain masks were created from the transformed masks in template space.

#### 2.3.2. Nonlinear template construction

As we have stated previously, it is desirable that a template not be biased towards any particular individual (or subset of individuals) in the population from which it is constructed. By biased, we mean that the template should not, on average, be more like any one subject than any other. There are two ways in which a template might appear more similar to an individual: in its shape, and in its appearance - where appearance refers to the voxel intensities. Consequently, a template may exhibit either a shape bias, an appearance bias, or both, unless care is taken to avoid this.

Shape (spatial) bias can be avoided by ensuring that, following registration to the template, the average displacement from the template to each individual is minimised across the population. Appearance (intensity) bias can be avoided by ensuring that, following registration to the template, the average image dissimilarity metric used to drive the registration is minimised across the population. Dissimilarity metrics commonly used by registrations methods include mean squared difference, cross-correlation and mutual information.

Fonov et al. (2011) formalised this concept as finding the template *T* that simultaneously minimises Equations 2 and 3, which address spatial and intensity bias respectively. The former (Eq. 2) minimises the magnitude of the nonlinear deformations 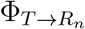 required to warp the template *T* to each subject *R*_*n*_, and the latter (Eq. 3) minimises the mean squared intensity difference between the template *T* and each warped subject 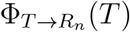. Note that Equation 3 is specific to our case where MMORF optimises an image dissimilarity metric that is a version of the sum of squared-differences, and would differ if, for example, cross-correlation was used instead.

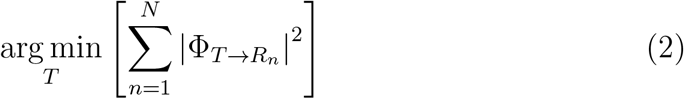

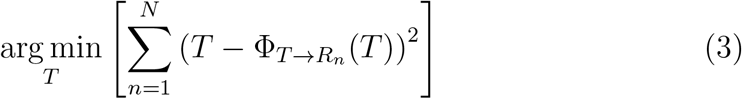

In practice, these two steps are interleaved at each of multiple iterations. In iteration *k*, Equation 2 is minimised by “undoing” (inverting and applying) the average across all nonlinear deformations 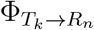 required to warp the template *T*_*k*_ to each subject *R*_*n*_, and Equation 3 is minimised by simple voxelwise averaging of the warped intensities 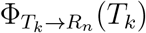 across all subjects. Given this understanding of unbiasing at each stage/iteration of the template construction pipeline, the optimal, unbiased, nonlinear OMM-1 template *T* was constructed by iterating over the following three steps (Fig. 1B).

1. Deformation fields 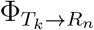 are estimated by nonlinearly registering each individual to the template from the previous iteration *T*_*k−*1_, with the affine template *T*_0_ being used as reference space for the first iteration. Registrations were performed with MMORF (Lange et al., 2020a) and were informed with both scalar modalities (T1, T2-FLAIR) and the tensor-valued modality (DTI) from individual and reference space. MMORF optimizes the following total cost function

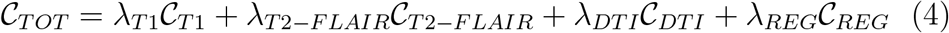

with all modalities contributing equally in the optimisation procedure (*i*.*e*., *λ*_*T*1_ = *λ*_*T*2*−F LAIR*_ = *λ*_*DT I*_ = 1). The mean squared error is calculated between scalar images and the mean squared Frobenius norm is used as a cost function for tensors. Registrations were initialised with the corresponding linear transformations *A*_*n*_ estimated during the construction of the affine template. Note that non-brain-extracted individual images and templates were used for the scalar channels, which poses additional challenges. Inclusion of the skull can negatively affect registration quality in nearby cortical regions, and the face and neck have larger anatomical and positional variability compared to the brain. To reduce the potential impact of extracranial tissue on the deformations close to the brain, and improve registration quality in the face and neck, different levels of relative regularisation were imposed on intra- and extracranial regions. This was achieved through modulation of the T1 brain mask in template space. Larger weights were given to regions inside the brain, i.e., reducing the relative level of regularisation to allow for more aggressive deformations, and smaller weights were given to regions outside the brain, i.e., increasing the relative level of regularisation to constrain the deformations. The intra-to-extracranial weight ratio was approximately 8-to-1. Similarly, a weighted average DTI mask with smoothly decreasing weights at the edge of brain tissue in reference space and eroded DTI brain masks in individuals’ native spaces were used to reduce the potential negative effect of poor/noisy tensor fitting around brain boundaries that are often seen in DTI. No masks were required for T2-FLAIR since tissue outside the brain already appears dark and does not strongly drive the registration relative to brain tissue.
2. The average deformation field 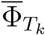 was calculated with

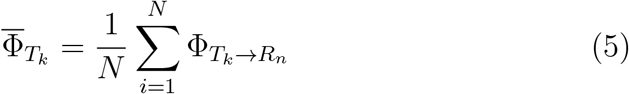

and used to spatially unbias the template. This unbiasing step was performed by composing its inverse 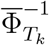 with each individual deformation field and the corresponding rigid and affine transformations:

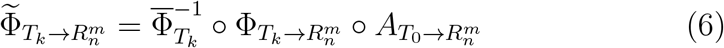 Individuals’ modalities in their respective native spaces were resampled to the new unbiased template space in one step by applying 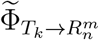 with spline interpolation. Tensors were reoriented with FSL’s *vecreg* tool, which uses the preservation of principal directions algorithm (Alexander et al., 2001). T1 and DTI brain masks were resampled with trilinear interpolation using the same transformations.
3. New mean masks and a new template *T*_*k*_ were created in unbiased space by taking the average over the resampled images for each modality. This new unbiased template with its three volumes served as a reference space in the next iteration *k* + 1. We performed a total of *K* = 18 iterations, allowing for coarse to fine improvements, with three iterations at each of six hierarchical levels. A large grid spacing of 32 mm and a blurring kernel of 8 mm FWHM was used for the MMORF registrations (step 1) in the three iterations at the first hierarchical level. These parameter values were halved for each level, down to 1 mm and 0.25 mm (respectively) at the last hierarchical level. An overview of the MMORF registration parameters can be found in Appendix B.

#### 2.3.3. Age-dependent template construction

UKB individuals *S* = {*S*_*j*_*}*_*j*=*1*…*J*_ (*J* = 37, 330) from the 45-82 year age range were first affinely and then nonlinearly registered to OMM-1. Similar to the previous nonlinear registrations, we used MMORF with all three MRI modalities (Fig. 1C) and the registration parameters in Appendix B. The estimated set of deformation fields 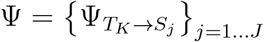 in OMM-1 space was used to model the average change in morphology with age using Gaussian process (GP) regression (Rasmussen and Williams, 2005), i.e. the objective can be stated as finding the function that best models the change in brain morphology as captured by the deformation fields given the subjects’ ages **x**. The trained GP allowed the prediction of a mean output deformation field 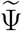 in OMM-1 space for any (observed or unobserved) age *x*_*\**_. Note that here we did not model differences in overall brain size and, consequently, the nonlinear deformation fields without their affine transformation components were used.

Gaussian processes generalize the concept of Gaussian probability distributions from stochastic variables to stochastic functions, and can be written as **f** (**x**) *∼GP* (*m*(**x**), *k*(**x, x**′)) or *y* = *f* (*x*) + *ϵ* with additive independent Gaussian noise *ϵ*. The GP is specified by its prior mean function *m*(**x**), which is usually set to zero, and covariance function *k*(**x, x**′), whose form has to be manually chosen. Conceptually these can be seen as continuous generalisations of the mean vector and covariance matrix used to describe multivariate normal distributions of random variables. The joint distribution of the observed training input and output pair (**x, y**) and unobserved pair (*x*_*\**_, *y*_*\**_) can be written as

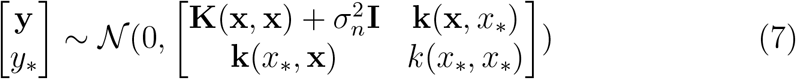

where **I** is the identity matrix and *σ*_*n*_ describes the variance of the noise, with larger *σ*_*n*_ resulting in a smoother function. **K**(**x, x**) is a *J* × *J* matrix of covariances between all training inputs, **k**(**x**, *x*_*\**_) and **k**(*x*_*\**_, **x**) are vectors of covariances between training and query inputs, and *k*(*x*_*\**_, *x*_*\**_) is the variance of the query input.

As will become more apparent from Equations 10-11, the calculation of 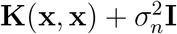 becomes increasingly computationally challenging with larger *J*. To reduce the computational burden, the training input and output data were stratified along the age axis into half-yearly bins. Additionally, each bin was split into two sub-bins, where each individual within an age bin was randomly assigned to one of the corresponding sub-bins. This introduction of some variability within each bin was done to better condition the estimation of the hyperparameters by making the estimates less correlated. This aggregation considerably reduced the size of the training dataset from the initial *J* = 37, 330 data points to *L* = 148 (2 sub-bins 74 half-yearly mean age bins 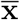) and corresponding mean deformation fields 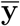 as respective input and output for ages 45-81 years.

The noise term in Equation 7 assumes that the noise is constant for every data point. This assumption would hold if every sub-bin was assigned the same number of individuals. However, when stratifying over age, the sub-bin averages were taken over different numbers of individuals because of the non-uniform age distribution in the initial dataset—with fewer individuals for the youngest and oldest age groups. Assuming noise to be constant for all subbins would introduce a bias. To account for this non-uniformity, the identity matrix **I** in Equation 7 was replaced with a weight matrix **W** containing 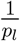 in the diagonal, where *p*_*l*_ is the number of individuals assigned to age bin *ℓ*. This down-weighs the noise variance for, and increases the confidence in, bins pooled from a larger number of individuals, and vice versa. The joint distribution from Equation 7 becomes

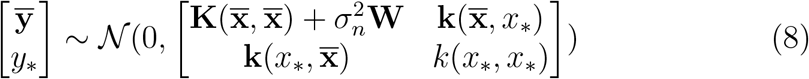

The choice of covariance function and its associated hyperparameters is important since it defines the properties of the functions generated during inference. Here, a squared exponential kernel was used, which has strong smoothness assumptions, and is therefore in line with the expected smooth changes in the brain with age. The kernel for calculating the covariance between two ages *x* and *x*′ can be written as

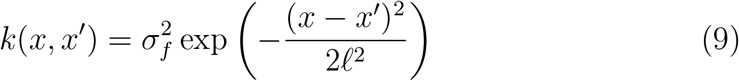

where *σ*_*f*_ is a scaling factor, and *ℓ* is the length scale. Intuitively, a larger *σ*_*f*_ increases both the magnitude and the variability of the fitted function, and a larger length scale *ℓ* increases the dispersion and covariance between more distant ages, leading to a smoother function, which is less influenced by noise and overfitting.

The hyperparameters *σ*_*f*_, *σ*_*n*_ and *ℓ* can be estimated by maximizing the marginal likelihood given by

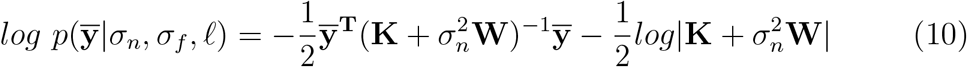

where 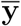is a matrix containing the vectorized deformation fields in the rows, and **K** is the covariance matrix where element *K*_*ij*_ is the covariance between two ages 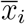and 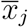. The Nelder-Mead simplex method has shown robust estimates when minimizing the negated function, which was optimized over all voxels to estimate the set of hyperparameters.

Using the estimated hyperparameters, the predictive mean can be calculated with

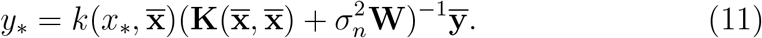

where the output *y*_*\**_ is the mean deformation field 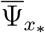 for the corresponding age *x*_*\**_. One deformation field was predicted for each year in the age range 45-81. The inverse of this deformation was concatenated with the initial 240 subjects’ linear and nonlinear transformations such that

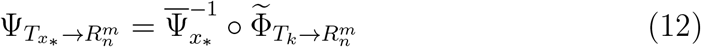

before resampling the corresponding modalities to age-specific template space in one step. Averaging over each of the resampled modalities provided the corresponding ADT 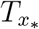.

### 2.4. Validation and applications

Convergence of the OMM-1 template construction process was assessed with three metrics including the root mean squared (RMS) difference, the root mean squared percentage (RMSP) difference, and Pearson’s correlation (PC) between consecutive iterations of average warps and T1, T2-FLAIR and DTI volumes. The Frobenius norm (FN) between consecutive iterations was additionally calculated for DTI volumes. Metrics based on the average warps show the improvement with respect to the first objective function of finding the average shape template, while metrics based on the T1, T2-FLAIR and DTI volumes show changes with respect to the second objective function of finding the average intensity template.

Our ADTs were visually assessed and the prediction-derived 81-year ADT was compared to two directly-estimated templates with the same age. The first of these two templates was constructed by registering all 101 UKB individuals in the 80-81 year range (mean age of 80.44 years) to OMM-1 space. The estimated deformation fields were spatially unbiased and applied to the corresponding images, which allowed the construction of a directly-estimated ADT-81. As a second template for comparison we used the existing older adult MIITRA template (mean age of 80.56 years) (Wu et al., 2022).

Age-related differences were assessed with the distortion given by average Jacobian determinant maps. One map was derived for each corresponding GP-predicted deformation field. Volume differences in the subcortical structures were quantified by summing over corresponding ROIs in the Jacobian determinant maps. The ROI masks were created by warping FSL FIRST (Patenaude et al., 2011) segmentation masks from individuals to OMM-1 space before averaging and binarizing them with a threshold of 0.5.

Finally, we investigated whether there is an advantage in registering individuals to the standard OMM-1 template via an age-matched ADT over registering them directly to the standard OMM-1. These results were also compared to registering directly to the MNI 152 template. 148 held-out UKB test subjects (50% female) were uniformly sampled from the 45-81 age range and registered to (1) the OMM-1 directly, (2) the GP-derived ADT corresponding to the individual’s age, and (3) the MNI 152 directly. FSL FLIRT (Jenkinson and Smith, 2001) was used for all affine registrations followed by MMORF (Lange et al., 2020a) for all nonlinear registrations. Registration parameters were identical for (1) and (2) with T1, T2-FLAIR and DTI driving the nonlinear registrations, and T1-only driving the nonlinear registration for (3). DKT atlas ROIs (Desikan et al., 2006) had been created for each individual with FreeSurfer (Fischl et al., 2004) as part of the UKB preprocessing pipeline. These ROIs were transformed from individuals’ native spaces to generic template space using the direct warp to OMM-1 as estimated in (1), the composed warp from individual to ADT and from ADT to OMM-1 for (2), and the direct warp to MNI 152 in (3) using trilinear interpolation. The Dice similarity coefficient for each transformed binarized ROI was calculated for every possible pairing of subjects for each of the three approaches. We repeated the same tests with 100 out-of-sample individuals and their corresponding Destrieux atlas Freesurfer ROIs (Destrieux et al., 2010) from the Human Connectome Project (HCP) (Van Essen et al., 2013) using T1-only, T1-only with brain extraction, and T1 and DTI modalities for registrations in (1) and (2). Due to the younger age range in the HCP, we used the youngest 45-year ADT as a reference space for all registrations in (2).

## 3. Results

### 3.1. OMM-1

The template after the last iteration of each hierarchical level is shown in Figure 2A. Qualitatively, a gradual increase in contrast and sharpness is noticeable in all three modalities with more rapid change over the early iterations and less change in the later iterations. Quantitative measurements of convergence towards the average intensity and average shape are shown in Figure 2B. The change in intensities and the change in average warps between consecutive iterations show a similar pattern with all metrics. A sharp increase in difference occurs after switching to a finer registration level before stabilising in the following iterations at the same level. These can be seen as sharp spikes in the differences measured with RMS, RMSP and FN, and sharp drops with PC. These fluctuations become smaller in the later iterations, which indicates convergence.

**Figure 2.**
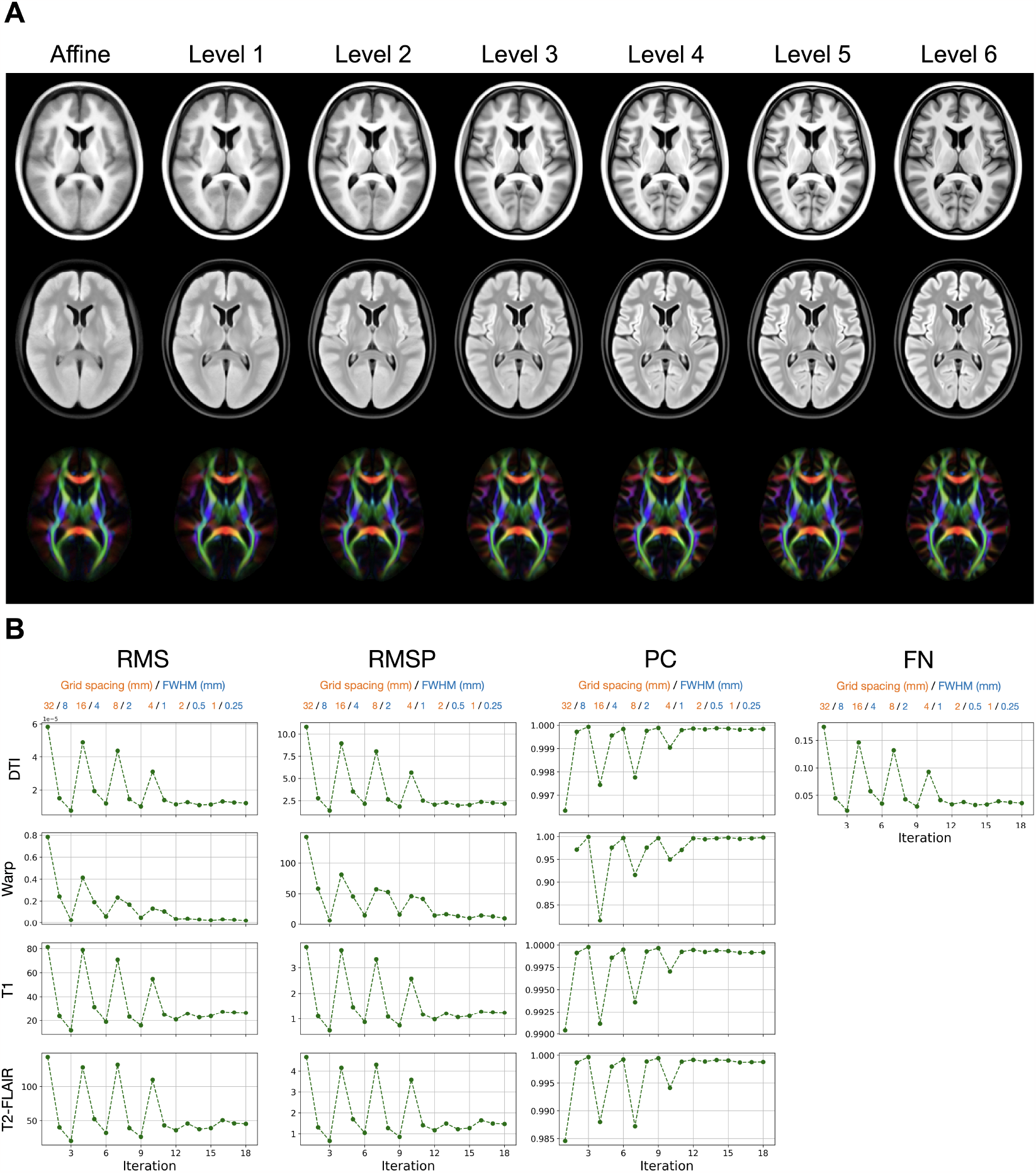
(A) Rows show the three template modalities: T1, T2-FLAIR and DTI (visualised using a principal diffusion direction colour-coded FA map where AP=Green, LR=Red, IS=Blue). The improvement in contrast and alignment after the final iteration of each hierarchical level is noticeable for each of the three modalities as the size of the blurring kernel and the grid spacing are reduced from coarse to fine. (B) Convergence measured with several metrics — root mean squared (RMS), root mean squared percentage (RMSP), Pearson correlation (PC) and Frobenius norm (FN) — shows the difference between template intensities for each modality and between average warps of consecutive iterations. A sharp difference can be observed with all metrics after switching to a new hierarchical level, which is followed by an improvement in the following iterations at the same level. These fluctuations stabilise towards the last iterations indicating convergence.

Two different sets of slices of the final OMM-1 template are shown in Figure 3. The T1 and T2-FLAIR volumes are visually sharp with excellent contrast. In the DTI volume, white matter fibre tracts appear clear and bright, and both isotropic and anisotropic regions are very well aligned. This is especially noticeable in the cerebellum and the brainstem tracts. All volumes, scalar, and DTI, show very good alignment, as judged by eye. This indicates that there are no systematic, rigid registration errors between modalities. Our strategy estimates a single warp from and applies it to all modalities in individual spaces, which ensures correct cross-modality alignment in template space (assuming that modalities in individual spaces are correctly aligned). Extracerebral structures, such as the sinuses are clearly visible in the T2-FLAIR volume, as well as the olfactory tract and the ol-factory bulbs. Subcortical GM structures are sharply defined, with complementary contrast provided by the T1 and T2-FLAIR modalities. (Fig. 3B).

**Figure 3.**
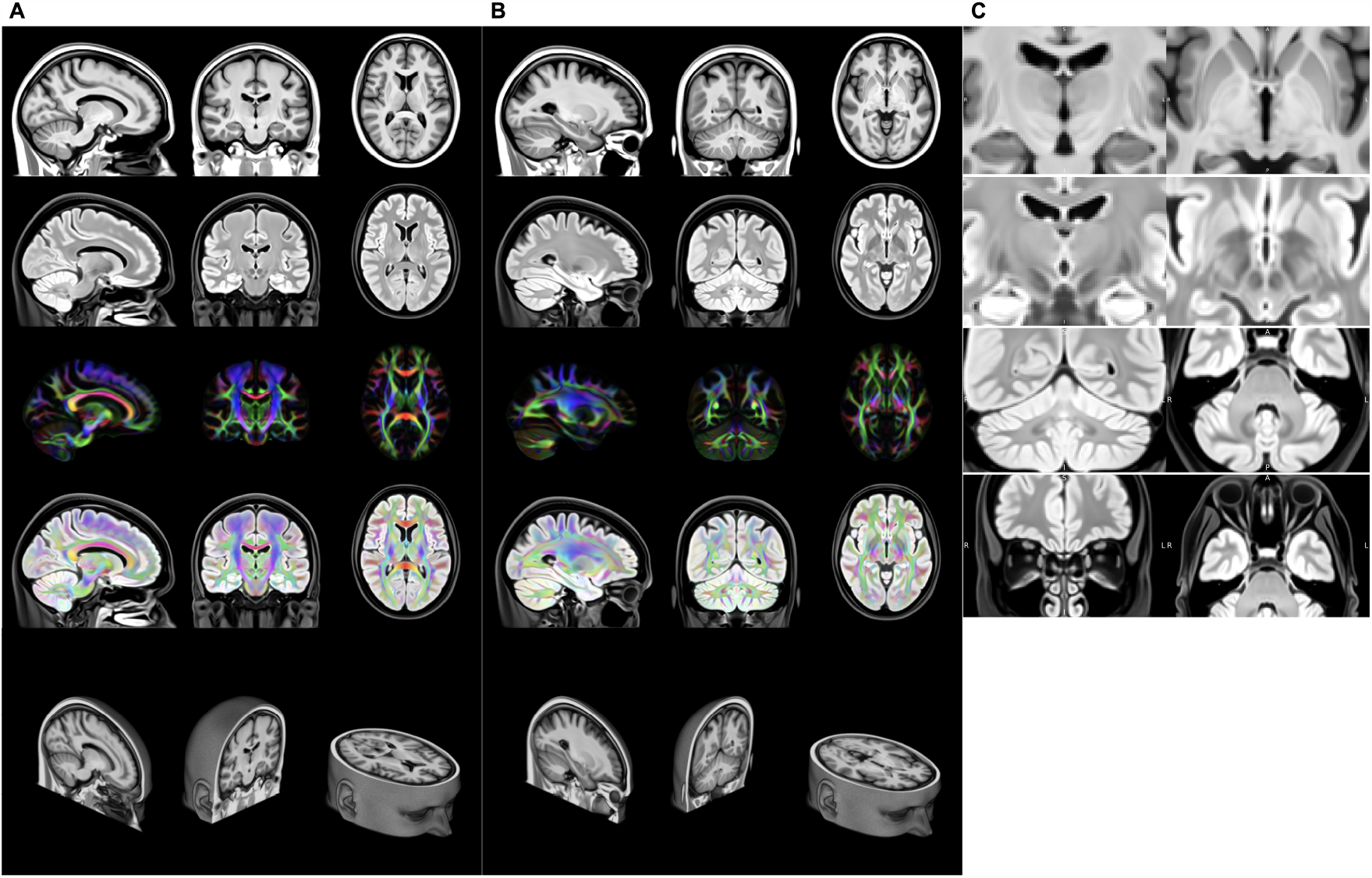
Two different sets of slices (A) and (B) through the modalities of the final OMM-1 template: T1, T2-FLAIR, DTI (visualised using a principal diffusion direction colour-coded FA map where AP=Green, LR=Red, IS=Blue), DTI overlaid on T2-FLAIR, and the corresponding slice in the 3D T1 volume. Scalar T1 and T2-FLAIR volumes exhibit very good contrast. The DTI volume shows excellent sharpness and orientational consistency. Alignment between scalar and tensor modalities is exceptional as can be seen in the overlay of DTI and T2-FLAIR. Facial features such as the ears, nose and eyes show a high level of detail. Coronal and axial slices show the right hemisphere on the left and vice versa. Zoomed-in views of four ROIs in (C) highlight the excellent alignment across all 240 participants. The medial medullary lamina - separating internal from external globus pallidus - is clearly visible in the axial view of the T1 volume. On the T2-FLAIR volume a clear separation of the subthalamic nucleus and substantia nigra can be seen in the coronal view, as well as the dentate nucleus in the cerebellum and the olfactory bulbs are clearly visible.

### 3.2. Age-dependent templates

The prediction-based ADTs for selected ages in steps of five years are illustrated in Figure 4A. The scalar and tensor volumes show consistently high quality of both contrast and crispness, and good alignment between modalities. Expected age-related differences, such as increases in ventricle sizes and sulcal widening, are visible in the templates, while the overall shape of anatomical brain structures and the folding pattern remains stable across all ages. This is confirmed when looking at the Jacobian determinant maps, where the largest differences occur in the ventricles. Cortical GM thinning is noticeable in the insular cortex and the inferior frontal gyrus.

**Figure 4.**
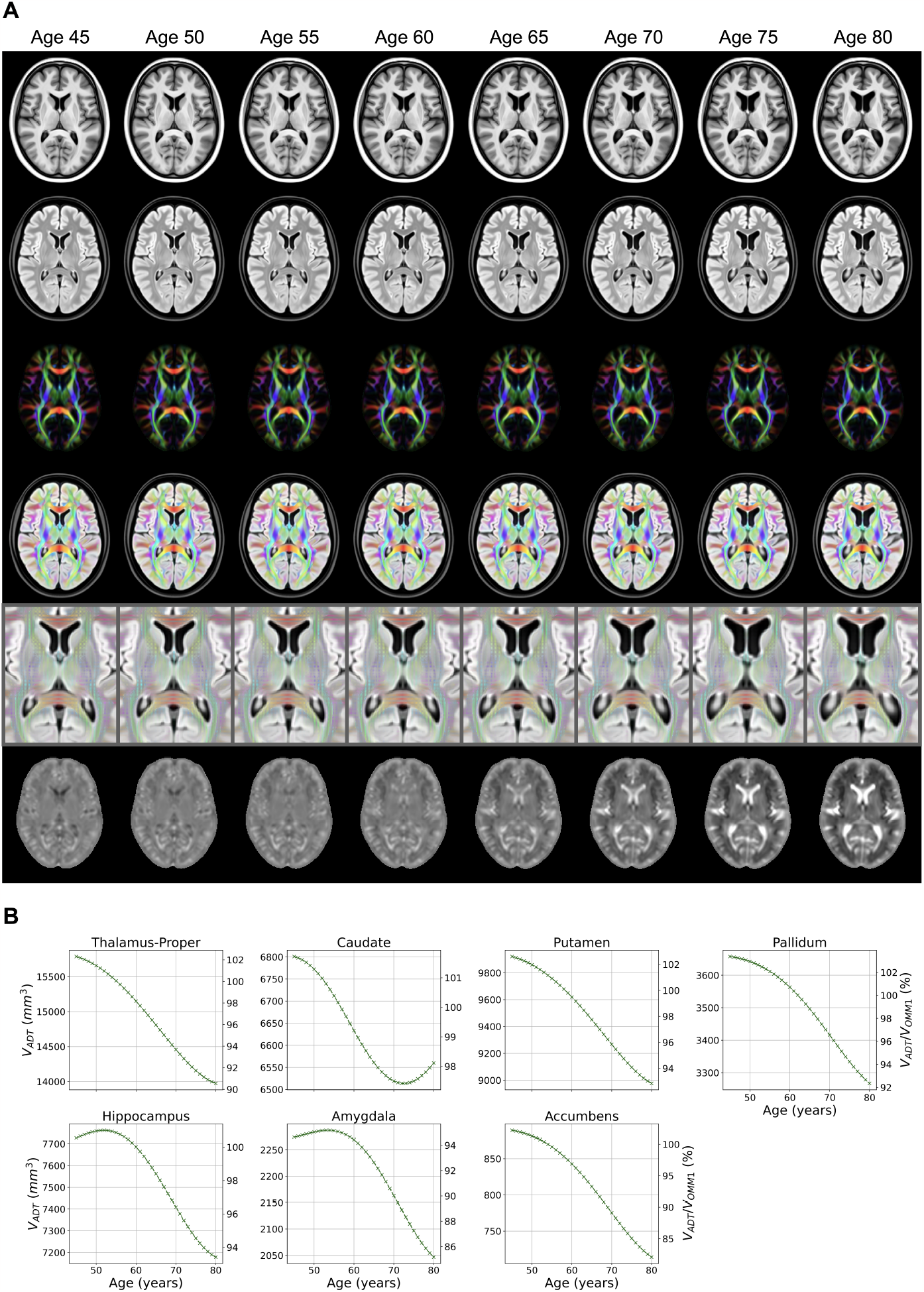
(A) The increase in ventricle size and sulcal widening can be observed across all modalities in this subset of age-dependent templates (right hemisphere on the left). The expansion seen in ventricles and sulci and the thinning of GM are visible in the corresponding log-Jacobian maps with dark values indicating contraction and bright values expansion of the main OMM-1 template. The *log*-|*J*| intensity range is set to *log*(0.5) *−log*(2.0). (B) Volume measurements directly derived from the templates show characteristic age-related differences in s ubcortical structures. Absolute volumes are given on the left y-axis and percent volume in comparison to OMM-1 on the right y-axis.

Age-related differences in volume can also be directly derived from the templates as exemplified in subcortical structures in Figure 4B, where an agerelated loss in volume is noticeable for all structures. A small unexpected increase in the hippocampus and amygdala volumes was found before age 52. We further investigated whether this increase is caused by an artefact of the modelling or the data by comparing measurements from the GP-estimated 48 year Jacobian determinant map with those derived from the average Jacobian determinant map of all 45-50 year old individuals. We found that our model underestimated the relative volume by approximately 1.5% for the hippocampus and 1.1 % for the amygdala in this age range, which will be partly caused by the smaller number of individuals in this age range, leading to larger uncertainty in the GP estimates. A similar pattern in the hippocampus was observed outside the main data range in (Janahi et al., 2022), where GPs were directly applied to volume measurements. However, for the hippocampus, similar trajectories in volume difference due to the data have been reported in the majority subgroup of the study population in (Fraser et al., 2021).

The increase in volume seen in the caudate from 72 years is an artefact caused by the segmentation masks, where the increasingly large ventricles start bleeding into the caudate ROI at older ages.

Figure 5 visually illustrates the age-related differences between the OMM-1 and the 81 year ADTs. It also highlights the high similarity between the directly-estimated ADT, the GP-estimated ADT, and the MIITRA template (Wu et al., 2022). Note that the GP-estimated ADT was derived by transforming the original 240 OMM-1 individuals from the 50-55 year age range through a GP-predicted warp, while the directly-estimated ADT was derived by transforming and averaging individuals with a mean age of 80.44 years. MIITRA was created through an iterative process from a cohort with a mean age of 80.56 years. While the general shape of all three templates is very similar, there are some differences. The directly-estimated ADT has slightly larger ventricles than both the GP-estimated ADT and MIITRA. It is also slightly blurrier in cortical areas compared to the GP-estimated ADT. Our GP-estimated ADT shows improved sharpness in subcortical areas, while MIITRA shows improved sharpness in cortical areas such as the occipital lobe. The striped appearance of the striatum is visible in the MIITRA, but not in the OMM-1 template.

**Figure 5.**
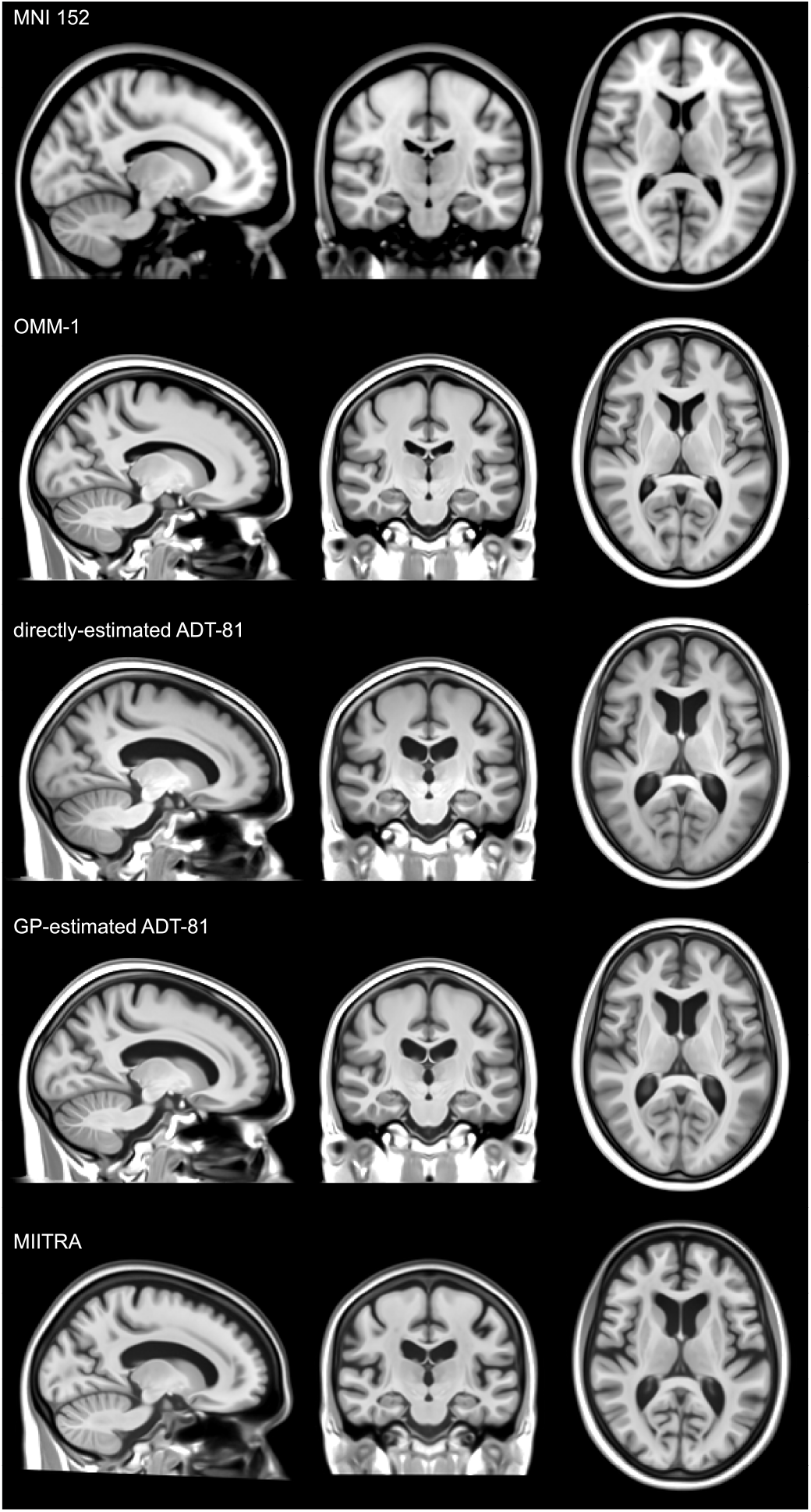
Visual comparison of the T1 volumes of the MNI 152, OMM-1, 81-year ADT directly estimated from 81 year old UKB individuals in OMM-1 space, 81-year ADT estimated with the GP, and MIITRA template (rigidly aligned) (Wu et al., 2022). The MNI 152 is provided as a reference and highlights the large global scale differences in MNI templates. Age-related difference between the OMM-1 and the two versions of the ADT-81 are well noticeable. The GP-estimated ADT-81 shows high similarity in overall shape and appearance, and age-related features such as ventricle size and sulcal widening, with both the directly-estimated ADT-81 and MIITRA. The GP-estimated ADT was derived from the GP-transformed, original 240 OMM-1 individuals (50-55 year age range). The directly-estimated ADT and MIITRA were derived from older adults with mean ages of 80.44 years and 80.56 years respectively.

The MNI 152 is provided as a commonly-used reference template and highlights the large difference in scale compared to the OMM-1, ADT and MIITRA templates. The cerebral volume of the OMM-1 is approximately 1,419,081 mm^3^, which is more than 465 ml less than the 1,884,594 mm^3^ of the MNI 152 and much more similar to the median and mean volumes of 1,433,335 mm^3^ and 1,440,417 mm^3^, respectively, in the entire UKB.

### 3.3. Application to spatial normalisation

Relative differences in aggregated pairwise dice coefficients for each ROI on UKB data are shown in Figure 6. On average over all ROIs spatial alignment to the corresponding ADT slightly outperforms alignment to OMM-1 by 0.64% in cortical and 0.42% in subcortical ROIs. Alignment to OMM-1 outperforms MNI 152 by a larger margin of 12.48% in cortical and 4.36% in subcortical ROIs. ADTs compared to OMM-1, and OMM-1 compared to MNI 152 achieve significantly larger Dice overlaps in 49 and 51 out of 63 cortical ROIs respectively. ADTs compared to OMM-1, and OMM-1 compared to MNI 152 achieve significantly larger Dice overlaps in 22 and 23 out of 32 subcortical ROIs respectively.

**Figure 6.**
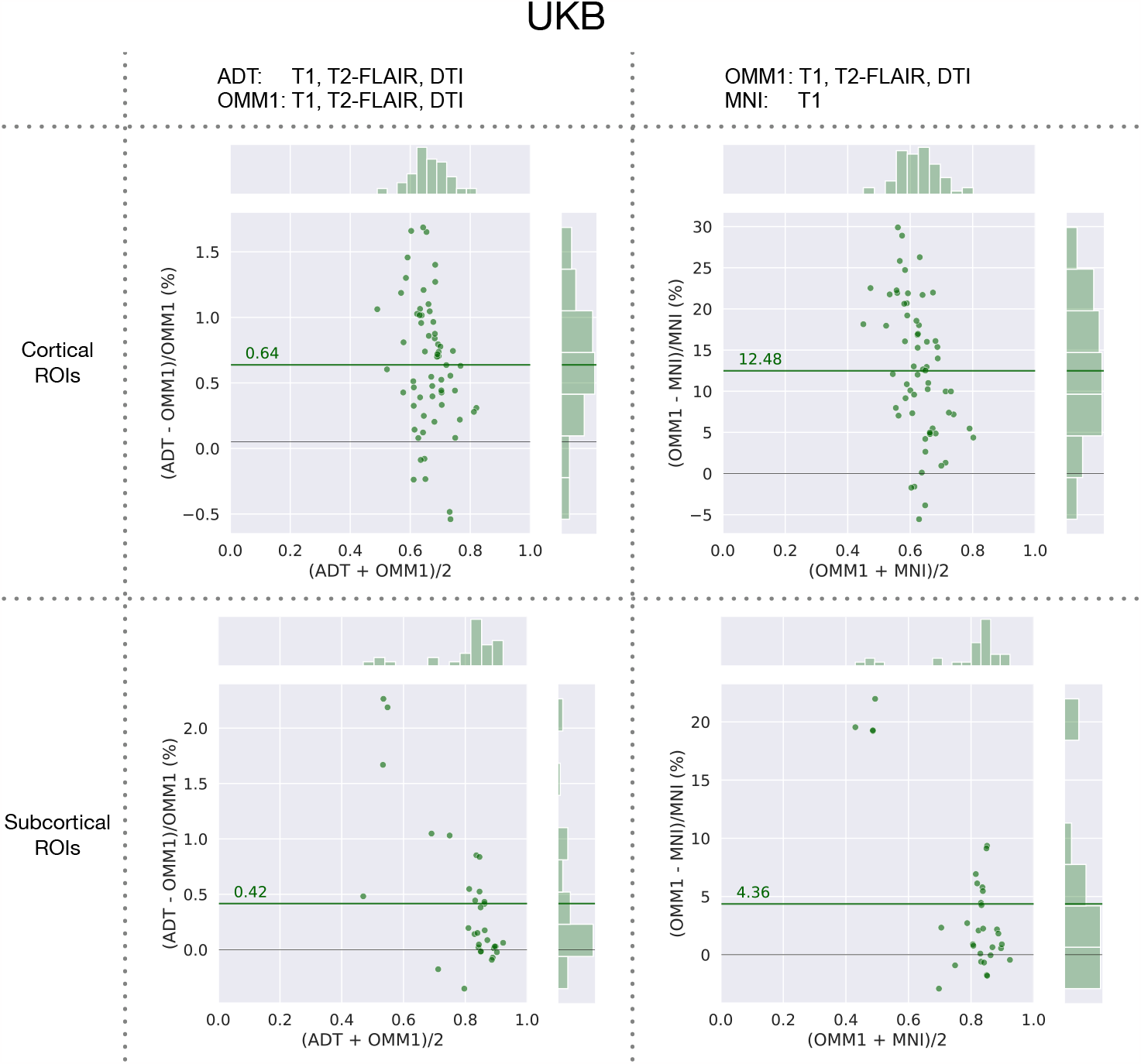
UK Biobank: relative differences between Dice coefficients obtained by warping the same masks with transformations estimated when directly registering to OMM-1 (OMM1), indirectly registering to OMM-1 via the individual’s corresponding age-dependent template (ADT), and registering to MNI 152. Each dot indicates one ROI and the green line shows the average percentage difference over all ROIs. On average ADTs slightly outperform OMM-1 and OMM-1 outperforms MNI by a larger margin.

Similar results can be replicated in an out-of-sample (non-UKB) cohort from the HCP (Fig. 7) where the 45 year ADT outperforms OMM-1 on average by 0.66% in cortical ROIs and 0.35% in subcortical ROIs. Multimodal registration to OMM-1 outperforms single-modal T1 registration to MNI by 3.31% and 1.77% respectively. Using only T1 for the registration to OMM-1 performs equally well as registration to MNI for cortical ROIs and 0.54% better for subcortical ROIs. Using preprocessed, brain-extracted images and templates for T1-based registrations showed a large improvement of OMM-1 over MNI 152 for both cortical and subcortical ROIs.

**Figure 7.**
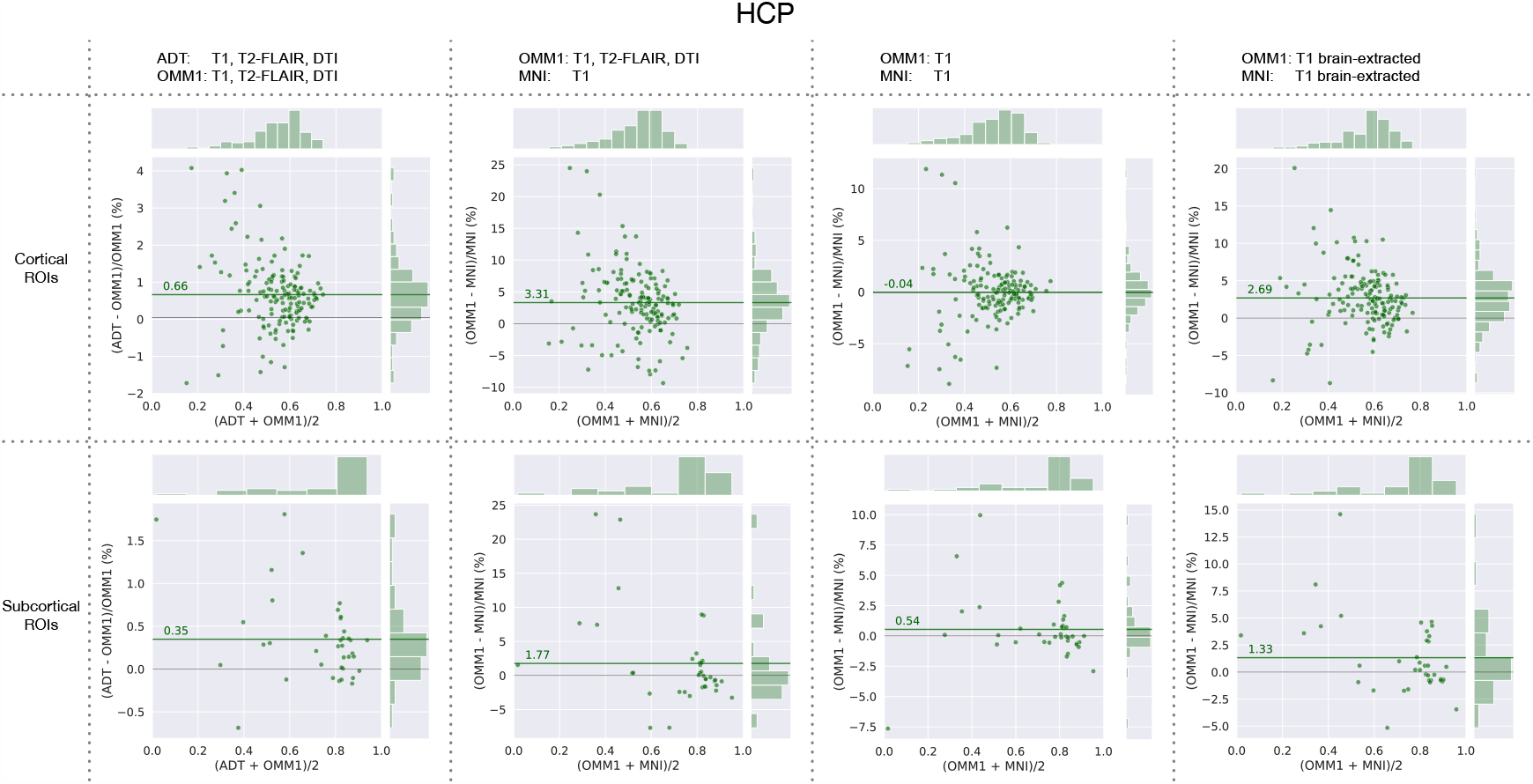
HCP: relative differences between Dice coefficients obtained by warping the same masks with transformations estimated when directly registering to OMM-1 (T1, DTI), indirectly registering to OMM-1 via the individual’s corresponding ADT, and registering to MNI 152. Additionally, registrations based on T1-only and brain-extraction T1-only were compared for OMM-1 and MNI 152. Each dot indicates one ROI and the green line shows the average percentage difference over all ROIs.

## 4. Discussion

We have presented the construction of the OMM-1, a fully-unbiased, internally-consistent, multimodal template of the brain including parts of the neck and face, averaging across 240 UKB individuals in the 50-55 year age range. GP regression was used to model deformation fields with age after registering 37,330 UKB individuals to OMM-1, and allowed the prediction of an average warp for each year of age in the corresponding 45-81 age range. These warp predictions were used to resample the original 240 OMM-1 subjects and create one multimodal ADT for each year of age. Test subjects from the UKB and HCP were registered to the OMM-1 directly, the ADT corresponding to the age of the individual, and MNI 152 to compare their performance in spatial normalisation tasks.

OMM-1 and ADTs consist of T1, T2-FLAIR and DTI volumes, and the same modalities were used to jointly inform the template construction process through the use of MMORF for all nonlinear registrations. The scalar volumes of all templates provide excellent contrast and exceptional anatomical detail, and the DTI volume appears sharp in isotropic and anisotropic regions. Our strategy inherently provides optimal cross-modality alignment between template volumes since all modalities are resampled through the same warps.

The OMM-1 is rigidly (6 DOF) aligned to MNI 152 space to provide a basic level of comparability, while avoiding scaling effects, to preserve the average brain size and shape of the UKB population. The scaling factor (Jacobian determinant of the affine transformation) between the OMM-1 and MNI 152 indicates an approximately 1.33 times larger cerebral volume of the MNI 152, which is also visually noticeable in Figure 5. Although this difference in scale might not have a large observable impact when used for spatial normalisation, it is not optimal as it will require additional unnecessary distortion for the majority of subjects. However, we recognise that considerable effort has been put into the development of atlases and analysis of studies in MNI space, and we therefore provide a deformation field that maps between MNI and OMM-1 space to allow these to be used with, or adapted for, our new template.

The GP model was used to create ADTs in steps of one year between ages 45 and 81, but allows for the construction of ADTs on a continuous scale within this age range. We did not test the prediction of templates outside this age range, since the GP model fit becomes increasingly uncertain when extrapolating.

Visual inspection (Figure 5) shows that our GP-estimated ADT-81 is highly similar in appearance to an ADT directly estimated from 81-year UKB individuals. It also has highly similar shape- and age-related features with respect to ventricle size, cortical folding and global shape and size compared to the MIITRA template, which was directly constructed from an older adult cohort. The slightly larger ventricles of the directly-estimated ADT will likely be caused by the smaller age range (80.0-81.0 years) of the subjects used in its construction. This is in stark contrast to 65.2-94.9 years for MIITRA and the weighted contributions according to the GP hyperparameters for the GP-estimated ADT.

The GP-estimated ADT shows improved sharpness in subcortical areas, while MIITRA shows improved sharpness in cortical areas, especially in the occipital lobe. Improvements in the MIITRA template in these areas are likely due to their weighted averaging approach, where intensities more similar to the median intensity across subjects at a voxel location receive higher weights. Notably, the striped appearance of the striatum is visible in the MIITRA, but not in the OMM-1. We have identified two reasonable causes for this. The first is that the regularisation metric used by MMORF to generate OMM-1, compared to that used by ANTs/DR-TAMAS to generate MIITRA, will more strongly penalise the deformations required to align the stripes across individuals. Since MMORF optimises the structural and DTI alignment simultaneously, the deformations required to align the stripes would negatively impact the alignment of the tensors in that region (particularly in terms of orientation alignment). This is supported by experiments (not shown) where we generated templates using only the T1 channel, and in which the stripes were partly visible (but still not to the extent seen in MIITRA). The second is that there does not appear to be a clear biological indication on the consistency of the stripes (= pontes grisei caudatolentiformes alternating with white matter forming the internal capsule) in number or exact location across individuals. Consequently, we do not believe that this negatively affects the use of OMM-1 as a registration target, even for older subjects.

We found that using the GP approach over simpler methods (such as kernel regression or the direct estimation of a template for each year) produced ADTs where morphology not affected by ageing (*e*.*g*., the folding pattern of the cortex) remained far more stable as a function of time. Our GP-based approach is also much more time-efficient compared to the repeated, direct construction of templates for specific ages. The most time-intensive tasks have to be performed only once, i.e., the iterative construction of a template (OMM-1) and the training of the GP. The estimated hyperparameters can then be re-used for the prediction of new templates thereafter.

The loss of subcortical volume in Figure 4 is mostly in line with results previously reported in the literature (Walhovd et al., 2005; Vinke et al., 2018; Nobis et al., 2019; Wang et al., 2019). The increase in hippocampus and amygdala volumes for the younger ages was unexpected and appears to be related to the smaller number of individuals available in this age range. This is similar to the results obtained by Janahi et al. (Janahi et al., 2022), where a GP was applied to extracted volume measurements. In comparison to direct measurements from the data, our model-derived ADTs underestimate the volume by approximately 1.1 % to 1.5 % for this age range. However, it should be noted that a similar increase in hippocampus volume for this age range has previously been reported (Fraser et al., 2021).

The increase in caudate volume seen from age 72 was caused by the increasing size of the ventricles at older ages bleeding into the caudate ROI. In addition to absolute estimated volumes, we show relative volumes normalized by the volumes estimated in generic OMM-1 space. These ratios change at different rates for different ROIs as would be expected. The volume change of the amygdala is a slight outlier in that it does not reach 100%. The reason for both of these deviations is the higher warp resolution of 1 mm used for the registrations in the construction of the OMM-1, compared to 2 mm for the GP-estimated ADTs. Increasing the warp resolution to 1 mm would produce more fine-grained average warps that might mitigate this effect. However, given the increasingly large size of the dataset and the corresponding number of required nonlinear registrations, 2 mm was found to be a reasonable compromise, and is comparable to, or better than, the standard settings of other registration methods.

The presence of pathologies such as white matter hyperintensities or microbleeds at older ages have an impact on registration and subsequently on segmentation accuracy in ADTs. Images of individuals in the 50-55 year age range provide enhanced tissue contrast, more detail in anatomical structures, and less pathologies than images from older individuals. This enhanced quality was the main motivation behind choosing 240 younger subjects for the construction of the OMM-1, and is an advantage when using the templates for spatial normalization, which is generally the main use case for population-based templates. Common age-related pathologies not present in these 240 individuals will also not be present in ADTs, whose image intensities are all derived from the the same 240 individuals. Similarly, DTI volumes of the ADTs will show expected differences in shape but not in the derived FA and MD maps. However, despite the use of these 240 younger individuals for the GP-estimated ADTs, appearance did not show substantial differences compared to the directly-estimated ADT-81 (from 80-81 year old UKB individuals) and the older-adult MIITRA template.

On average, alignment to OMM-1 via registration to an individual’s age-corresponding ADT shows slightly better spatial normalization performance than registering directly to OMM-1 in both UKB and HCP test subjects. Although the MNI 152 template is outside the age range of the UKB, it is commonly used for adult studies of all ages and, as such, it was included in our comparison. The use of OMM-1 and ADTs as template spaces outperformed MNI 152 on both UKB and HCP test subjects for the majority of ROIs. The performance of MNI 152 was similar to T1-only registration to OMM-1 when the skull was included. This can be explained by the large scalp signal present in the UKB T1 images and, consequently, also in OMM-1 and ADTs that does not match the characteristics of HCP data. The sharp improvement of T1-only registrations when used with a brain mask can be observed in Figure 7. Hence, we recommend the use of a brain mask for the T1 template volume when used as registration reference in uni- and multimodal registrations. It is also recommended to use a mask for DTI to avoid any potential impact of noisy tensors outside the brain. We supply such masks in template space as part of our OMM-1 release.

Whether or not the benefits of slightly improved registration outweigh the added complexity of using ADTs will likely depend on the specifics of a particular study. In many cases it may be sufficient to simply use the OMM-1 directly. However, should the use of an ADT be preferred (e.g., the 80 year ADT for an older population study), then our template provides a natural way to compare results with those from other studies using the generic OMM-1, as well as making atlases defined in generic OMM-1 space available in any ADT. In addition to age, a new GP model could also be conditioned on other attributes such as sex. To get high confidence predictions, this approach requires datasets with sufficiently well-represented sub-populations with the attribute of interest.

Our multimodal templating strategy provides a framework for integrating complemental information from scalar- and tensor-valued modalities with MMORF in a fully-unbiased and internally-consistent way. The use of our templates in combination with MMORF can largely improve accuracy in spatial normalisation tasks and the availability of spatially corresponding information from anatomical images such as T1 and T2-FLAIR and diffusion tensors from dMRI will greatly benefit the interpretation of results in template space. OMM-1 and all pre-constructed ADTs, the code for template construction, and MMORF and the MMORF config files used for the non-linear registrations can be publicly accessed via (Arthofer et al., 2023). Our template construction pipeline is not limited to these specific modalities and can be readily applied to other modalities and datasets of interest. In the future we hope to further extend the field of view of the OMM-1 to include the whole neck and face, which could add further benefits for MEG and EEG studies.

## 5. Acknowledgements

This research was funded by a Wellcome Trust Collaborative Award (215573/Z/19/Z). The Wellcome Centre for Integrative Neuroimaging is supported by core funding from the Wellcome Trust (203139/Z/16/Z). Computation used the Oxford Biomedical Research Computing (BMRC) facility, a joint development between the Wellcome Centre for Human Genetics and the Big Data Institute supported by Health Data Research UK and the NIHR Oxford Biomedical Research Centre. The views expressed are those of the author(s) and not necessarily those of the NHS, the NIHR or the Department of Health. This research has been conducted using the UK Biobank Resource under Application Number 8107. Data were provided in part by the Human Connectome Project, WU-Minn Consortium (Principal Investigators: David Van Essen and Kamil Ugurbil; 1U54MH091657) funded by the 16 NIH Institutes and Centers that support the NIH Blueprint for Neuro-science Research; and by the McDonnell Center for Systems Neuroscience at Washington University.

## Appendix A. Dealing with high intensity values

Clamping of high scalp intensities was achieved with a custom sigmoid clamping function

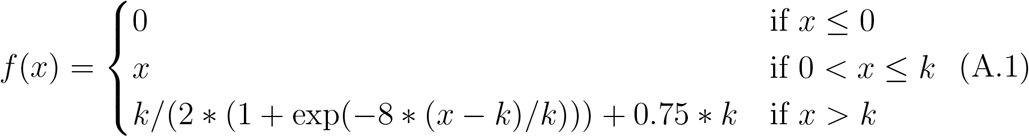

where *x* is a voxel’s intensity and *k* the mean WM intensity.

## Appendix B. MMORF registration parameters

Table Appendix B contains the MMORF registration parameters used for the construction of the OMM-1 and age-dependent templates. The main difference is in the maximum warp resolution which is 1 mm for the former and 2 mm for the latter. The choice of reducing the warp resolution from 1 mm to 2 mm was mainly motivated by savings in runtime, while still maintaining a high level of detail.

**Table B.3:** MMORF parameters used for the registrations when constructing the OMM-1 and age-dependent templates. Given the large number of individuals used in the age-dependent template modelling and the large associated computational requirements the highest warp resolution was set to 2 mm, in contrast to 1 mm used for the OMM-1.

|  | OMM-1 | Age-dependent templates |
| --- | --- | --- |
| ; general |  |  |
| warp_res_init | 32 | 32 |
| warp_scaling | 1 1 2 2 2 2 2 | 1 1 2 2 2 2 |
| lambda_reg | 4.0e5 3.7e-1 3.1e-1 2.6e-1 2.2e-1 1.8e-1 1.5e-1 | 4.0e5 3.7e-1 3.1e-1 2.6e-1 2.2e-1 1.8e-1 |
| hires | 3.9 | 3.9 |
| optimiser_max_it_lowres | 10 | 5 |
| optimiser_max_it_hires | 10 | 5 |
| ; whole head T1 |  |  |
| use_implicit_mask | 0 | 0 |
| use_mask_ref_scalar | 1 1 1 1 1 1 1 | 1 1 1 1 1 1 |
| use_mask_mov_scalar | 0 0 0 0 0 0 0 | 0 0 0 0 0 0 |
| fwhm_ref_scalar | 8.0 8.0 4.0 2.0 1.0 0.5 0.25 | 8.0 8.0 4.0 2.0 1.0 0.5 |
| fwhm_mov_scalar | 8.0 8.0 4.0 2.0 1.0 0.5 0.25 | 8.0 8.0 4.0 2.0 1.0 0.5 |
| lambda_scalar | 1 1 1 1 1 1 1 | 1 1 1 1 1 1 |
| estimate_bias | 1 | 1 |
| bias_res_init | 32 | 32 |
| lambda_bias_reg | 1e9 1e9 1e9 1e9 1e9 1e9 1e9 | 1e9 1e9 1e9 1e9 1e9 1e9 |
| ; whole head FLAIR |  |  |
| use_implicit_mask | 0 | 0 |
| use_mask_ref_scalar | 0 0 0 0 0 0 0 | 0 0 0 0 0 0 |
| use_mask_mov_scalar | 0 0 0 0 0 0 0 | 0 0 0 0 0 0 |
| fwhm_ref_scalar | 8.0 8.0 4.0 2.0 1.0 0.5 0.25 | 8.0 8.0 4.0 2.0 1.0 0.5 |
| fwhm_mov_scalar | 8.0 8.0 4.0 2.0 1.0 0.5 0.25 | 8.0 8.0 4.0 2.0 1.0 0.5 |
| lambda_scalar | 1 1 1 1 1 1 1 | 1 1 1 1 1 1 |
| estimate_bias | 1 | 1 |
| bias_res_init | 32 | 32 |
| lambda_bias_reg | 1e9 1e9 1e9 1e9 1e9 1e9 1e9 | 1e9 1e9 1e9 1e9 1e9 1e9 |
| ; DTI |  |  |
| use_mask_ref_tensor | 1 1 1 1 1 1 1 | 1 1 1 1 1 1 |
| use_mask_mov_tensor | 1 1 1 1 1 1 1 | 1 1 1 1 1 1 |
| fwhm_ref_tensor | 8.0 8.0 4.0 2.0 1.0 0.5 0.25 | 8.0 8.0 4.0 2.0 1.0 0.5 |
| fwhm_mov_tensor | 8.0 8.0 4.0 2.0 1.0 0.5 0.25 | 8.0 8.0 4.0 2.0 1.0 0.5 |
| lambda_tensor | 1 1 1 1 1 1 1 | 1 1 1 1 1 1 |

## Footnotes

1 Only the b=1000 shell is used for tensor fitting due to the violation of the Gaussian diffusion assumption underlying the diffusion tensor model at higher b-values (or when combining b-values), which would require Kurtosis for correct modelling.

## Notes

https://doi.org/10.17605/OSF.IO/S9GE4

### Competing Interest Statement

The authors have declared no competing interest.

